## Supplemental Data for "Heterochromatin epimutations impose mitochondrial dysfunction to confer antifungal resistance"

### Supplementary Figures & Tables

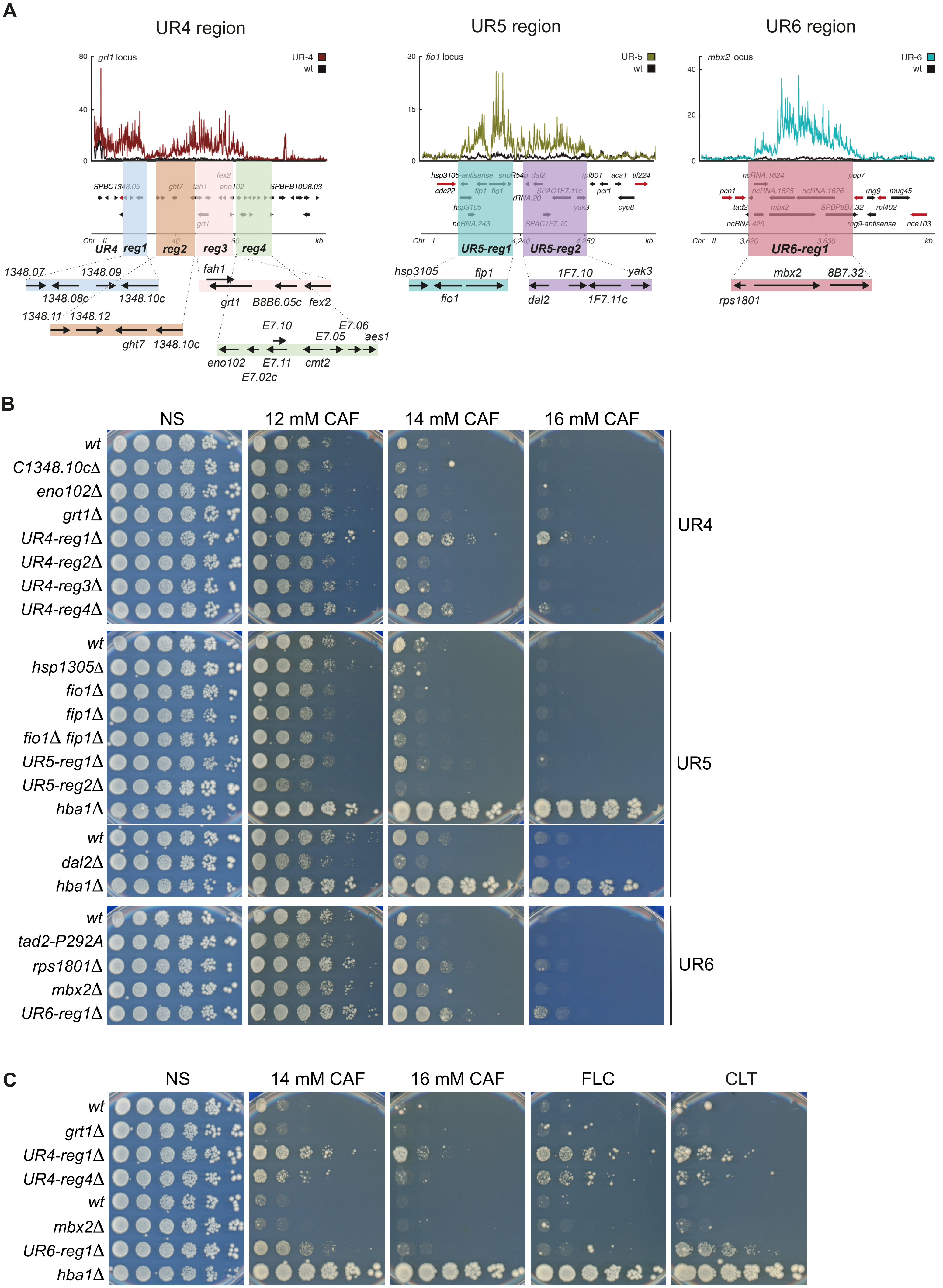

A
UR4 region
UR5 region
UR6 region
grt1 locus
fio1 locus
mbx2 locus
UR-4
UR-5
UR-6
wt
UR4 reg1 reg2 reg3 reg4
UR5-reg1 UR5-reg2
UR6-reg1
1348.07 1348.09 1348.08c 1348.10c
1348.11 1348.12 ght7 1348.10c
fah1 grt1 B8B6.05c fex2
E7.10 E7.06 E7.05 aes1 eno102 E7.11 cmt2 E7.02c
hsp3105 fip1 fio1
1F7.10 yak3 dal2 1F7.11c
mbx2 8B7.32 rps1801
B
NS
12 mM CAF
14 mM CAF
16 mM CAF
wt
C1348.10cΔ
eno102Δ
grt1Δ
UR4-reg1Δ
UR4-reg2Δ
UR4-reg3Δ
UR4-reg4Δ
UR4
wt
hsp1305Δ
fio1Δ
fip1Δ
fio1Δ fip1Δ
UR5-reg1Δ
UR5-reg2Δ
hba1Δ
wt
dal2Δ
hba1Δ
UR5
wt
tad2-P292A
rps1801Δ
mbx2Δ
UR6-reg1Δ
UR6
C
NS
14 mM CAF
16 mM CAF
FLC
CLT
wt
grt1Δ
UR4-reg1Δ
UR4-reg4Δ
wt
mbx2Δ
UR6-reg1Δ
hba1Δ

**Figure S1: Deletion of genes within regions where heterochromatin islands are located in UR4, UR5 and UR6 epimutants**

**A** Genes encompassed in heterochromatin islands present in UR4, UR5 and UR6 epimutants. Schematics of regions containing ectopic heterochromatin islands. Protein-coding genes only are indicated. Shaded blocks indicate the regions deleted in multiple-gene deletions, with expanded view below, where some genes are indicated with abbreviated names. *Left:* UR4 heterochromatin island, ChrII (0-60 kb); *centre*: UR5 heterochromatin island, ChrI (4230-4260 kb); right: UR6 heterochromatin island, ChrII (3615-3640 kb) ChIP-seq data is from (Torres-Garcia et al., 2020b).
**B** Growth assay to assess growth of deletion strains on caffeine. Single genes or regions containing multiple genes were deleted from wild-type cells and the ability of resultant strains to grow on media containing caffeine assessed. Five-fold serial dilutions of indicated strains spotted onto non-selective plates (NS; YES media) or plates containing the indicated concentrations of caffeine. Plates photographed after 2-8 days at 32$^{o}$C.
**C** Growth assay to assess resistance to caffeine and antifungal drugs. Strains which showed some resistance to caffeine in **B** were retested in growth assays to assess ability to grow on various insults. Five-fold serial dilutions of indicated strains spotted onto non-selective plates (NS; YES media) or plates containing 14 mM or 16 mM caffeine (CAF), 0.3 mM fluconazole (FLC), 50 ng/ml clotrimazole (CLT). Plates photographed after 2-8 days at 32$^{o}$C.

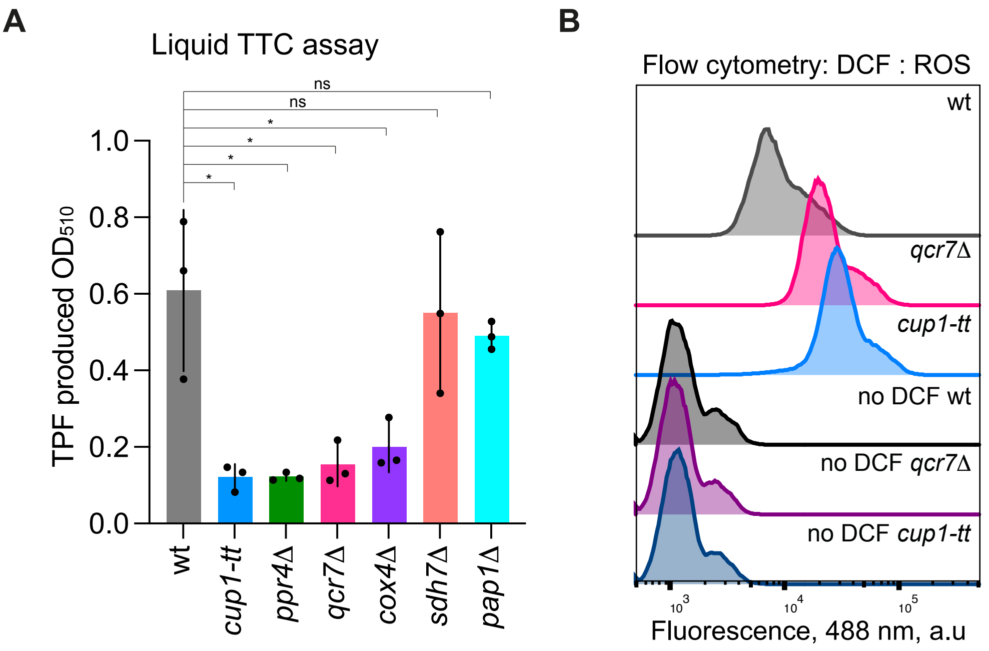

A
Liquid TTC assay
ns
ns
*
*
*
*
1.0
0.8
0.6
0.4
0.2
0.0
TPF produced OD510
wt
cup1-tt
ppr4Δ
qcr7Δ
cox4Δ
sdh7Δ
pap1Δ
B
Flow cytometry: DCF : ROS
wt
qcr7Δ
cup1-tt
no DCF wt
no DCF qcr7Δ
no DCF cup1-tt
10^3
10^4
10^5
Fluorescence, 488 nm, a.u

**Figure S2: Deficiency of Cup1, Ppr4 or ETC components cause respiratory deficiency**

**A** Liquid Tetrazolium assay for respiratory competence. Cells were incubated in 2,3,5-Triphenyltetrazolium Chloride (TTC) for 30 min before cell lysis and extraction with DMSO to release 1,3,5-triphenylformazan (TPF). TPF was measured by absorbance at 510 nm. Data are mean and standard deviation from three biological replicates. p values determined by two-tailed Student's t-test: * $p < 0.05$; ** $p < 0.01$; ns, not significant.

**B** Flow cytometry of cells stained with DCHF-DA or unstained wt, *qcr7*Δ and *cup1-tt* cells are shown.

**A**

| | *Source* | *Term ID* | *Term name* | *P-value adjusted* |
|---|---|---|---|---|
| *1* | GO:MF | GO:0016491 | oxidoreductase activity | 4.30006E-05 |
| *2* | GO:MF | GO:0004033 | aldo-keto reductase (NADP) activity | 0.003023053 |
| *3* | GO:BP | GO:0033212 | iron import into cell | 3.03281E-06 |
| *4* | GO:BP | GO:0098754 | detoxification | 0.003679474 |

| | *Source* | *Term ID* | *Term name* | *P-value adjusted* |
|---|---|---|---|---|
| *1* | KEGG | KEGG:00052 | Galactose metabolism | 0.004121623 |
| *2* | KEGG | KEGG:00750 | Vitamin B6 metabolism | 0.015561056 |

**B**

| | *Source* | *Term ID* | *Term name* | *P-value adjusted* |
|---|---|---|---|---|
| *1* | GO:MF | GO:0022857 | transmembrane transporter activity | 1.05E-12 |
| *2* | GO:MF | GO:0016491 | oxidoreductase activity | 2.00E-08 |
| *3* | GO:MF | GO:0016874 | ligase activity | 7.12E-07 |
| *4* | GO:MF | GO:0036094 | small molecule binding | 0.00048527 |
| *5* | GO:BP | GO:0009060 | aerobic respiration | 1.35E-17 |
| *6* | GO:BP | GO:0044281 | small molecule metabolic process | 2.06E-11 |
| *7* | GO:BP | GO:1902600 | proton transmembrane transport | 1.59E-09 |
| *8* | GO:BP | GO:0015711 | organic anion transport | 0.02213762 |
| *9* | GO:CC | GO:0005739 | mitochondrion | 1.74E-14 |

| | *Source* | *Term ID* | *Term name* | *P-value adjusted* |
|---|---|---|---|---|
| *1* | KEGG | KEGG:01100 | Metabolic pathways | 7.72E-14 |
| *2* | KEGG | KEGG:00190 | Oxidative phosphorylation | 1.35E-12 |
| *3* | KEGG | KEGG:00670 | One carbon pool by folate | 0.00092928 |
| *4* | KEGG | KEGG:01110 | Biosynthesis of secondary metabolites | 0.00418292 |
| *5* | KEGG | KEGG:01230 | Biosynthesis of amino acids | 0.0163563 |
| *6* | KEGG | KEGG:01210 | 2-Oxocarboxylic acid metabolism | 0.0388758 |
| *7* | KEGG | KEGG:01200 | Carbon metabolism | 0.04353672 |

**Figure S3: GO and KEGG analysis of *cup1-tt* and *ppr4*Δ transcriptomes**

**A** Enrichment analysis for overlapping induced genes in *ppr4Δ* and *cup1-tt*.

*Upper:* List of driver terms from the two-stage filtering algorithm in g:Profiler that systematically reduces the terms retrieved by incorporating information for the underlying GO-term topology. *Lower:* List of KEGG terms enriched. Link to full term list including nondriver terms in g:Profiler: https://biit.cs.ut.ee/gplink/l/xh7rrByhR2

**B** Enrichment analysis for overlapping repressed genes in *ppr4D* and *cup1-tt*.

*Upper:* List of driver terms from the two-stage filtering algorithm in g:Profiler that systematically reduces the terms retrieved by incorporating information for the underlying GO-term topology. Most DEGs in GO:MF transmembrane transporter activity map to mitochondrial genes. *Lower:* List of KEGG terms enriched. Link to full term list including nondriver terms in g:Profiler: https://biit.cs.ut.ee/gplink/l/Vl4bFVCkT6.

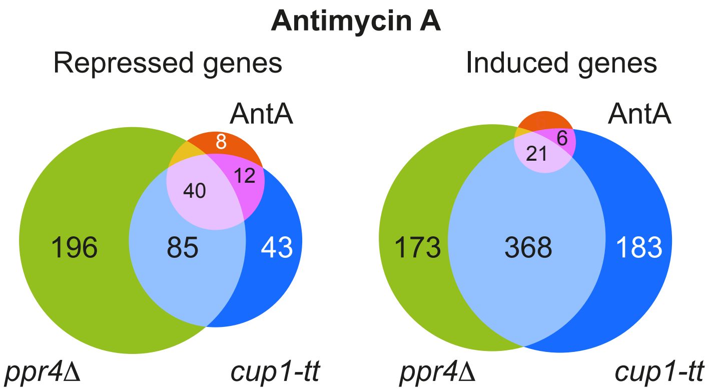

Antimycin A
Repressed genes
Induced genes
AntA
8
12
40
196
85
43
ppr4Δ
cup1-tt
AntA
6
21
173
368
183
ppr4Δ
cup1-tt

**Figure S4: Transcriptional profiles of cells defective for Cup1 or Ppr4 overlap with mitonuclear retrograde response**

Comparison of *cup1-tt* and *ppr4*∆ with antimycin A treatment (Malecki et al., 2016) *Left:* Venn diagram of comparison between genes repressed in *cup1-tt* (dark blue) and *ppr4*∆ (green) and downregulated in response to Antimycin A, which activates the mitonuclear retrograde pathway (pink). *Right:* Venn diagram of comparison between genes upregulated in *cup1-tt* (dark blue) and *ppr4*∆ (green) and induced in response to Antimycin A (pink).

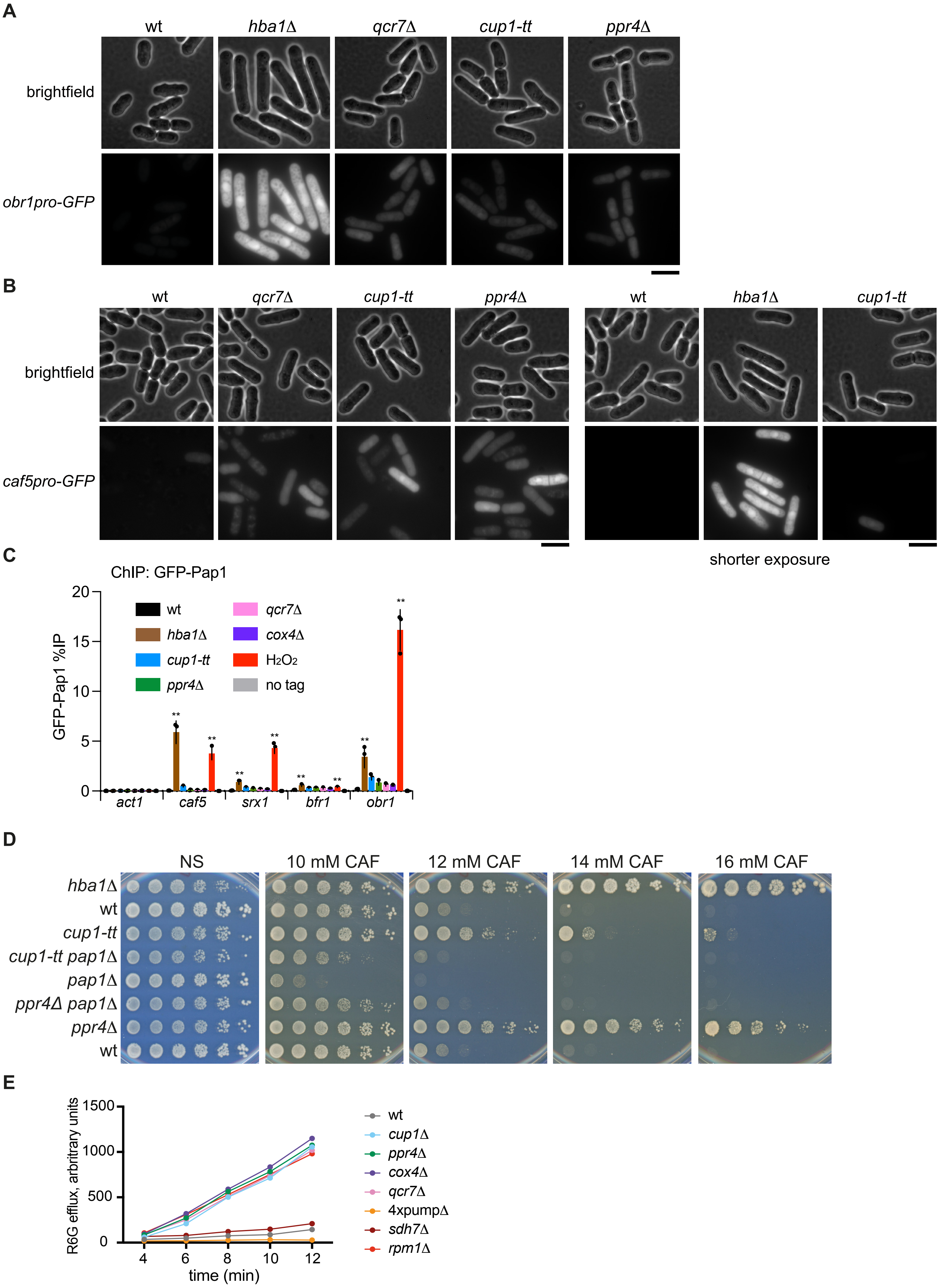
A
wt
hba1Δ
qcr7Δ
cup1-tt
ppr4Δ
brightfield
obr1pro-GFP
B
wt
qcr7Δ
cup1-tt
ppr4Δ
wt
hba1Δ
cup1-tt
brightfield
caf5pro-GFP
shorter exposure
C
ChIP: GFP-Pap1
GFP-Pap1 %IP
wt
hba1Δ
cup1-tt
ppr4Δ
qcr7Δ
cox4Δ
H2O2
no tag
act1
caf5
srx1
bfr1
obr1
D
NS
10 mM CAF
12 mM CAF
14 mM CAF
16 mM CAF
hba1Δ
wt
cup1-tt
cup1-tt pap1Δ
pap1Δ
ppr4Δ pap1Δ
ppr4Δ
wt
E
R6G efflux, arbitrary units
time (min)
wt
cup1Δ
ppr4Δ
cox4Δ
qcr7Δ
4xpumpΔ
sdh7Δ
rpm1Δ

**Figure S5: Cup1, Ppr4 and ETC deficient cells activate the Pap1-dependent stress response and show increased efflux**

**A** Pap1-dependent *obr1*-promoter reporter assay. Fluorescence microscopy images of live cells of indicated strains containing GFP under control of the Pap1-dependent *obr1* promoter. Fluorescence images are scaled relative to the brightest image (*hba1*Δ). Scale bar, 10 μm.

**B** Pap1-dependent *caf5* promoter reporter assay. Fluorescence and brightfield images live cells of indicated strains containing GFP under control of the Pap1-dependent *caf1* promoter. *Left panels*, 2000 ms exposure. *Right panels*, 300 ms exposure. Fluorescence images are scaled relative to the brightest image in each set of images. Scale bar, 10 μm.

**C** GFP-Pap1 ChIP-qPCR of the indicated strains and GFP-Pap1 wild-type cells treated with 0.2 mM hydrogen peroxide for 30 min. Promoter regions of the *caf5* and *bfr1* (transmembrane transporters), *srx1* (sulfiredoxin), and *obr1* (dehydrogenase) genes were analysed by qPCR to determine GFP-Pap1 % immunoprecipitation. *act1* serves as negative control locus. Data are mean and standard deviation from three biological replicates. p values determined by two-tailed Student's t-test: * p <0.05; ** p<0.01; ns, not significant. Only p values for wt vs *hba1*Δ and $H_2O_2$ treatment are indicated. p values for other mutants are shown in **Figure 5C**.

**D** Growth assay to assess impact of loss of Pap1 on *cup1-tt* and *ppr4*Δ mutants. Five-fold serial dilutions of indicated strains spotted onto non-selective plates (NS; YES media) or plates containing the indicated concentrations of caffeine. Plates photographed after 2-8 days at 32°C.

**E** Efflux of Rhodamine 6G (R6G) from cells. Cells of the indicated strains were preloaded with R6G and enabled to perform efflux when supplied with glucose (in YES). R6G released to the media over the indicated time-period was measured. Representative example shown. *4xpump*Δ strain four transmembrane transporters are absent (Bfr1, Pmp1, Caf5, Mfs1) along with Pap1 and Prt1.

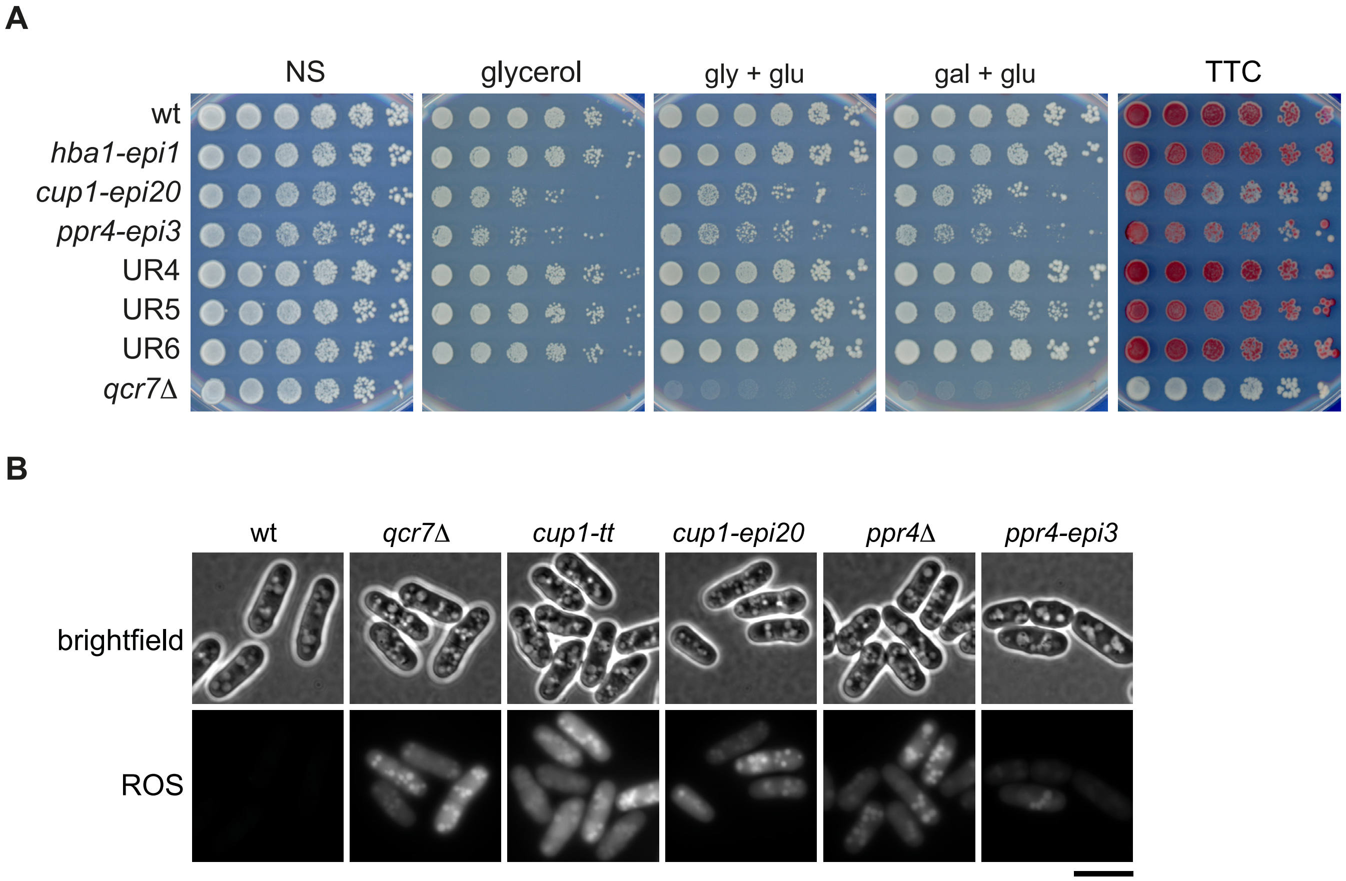
A
NS
glycerol
gly + glu
gal + glu
TTC
wt
hba1-epi1
cup1-epi20
ppr4-epi3
UR4
UR5
UR6
qcr7Δ
B
wt
qcr7Δ
cup1-tt
cup1-epi20
ppr4Δ
ppr4-epi3
brightfield
ROS

**Figure S6: Assessment of mitochondrial competence and ROS levels in epimutants**

**A** Growth assay to assess respiratory competence. Five-fold serial dilutions of indicated strains spotted onto non-selective plates (NS; YES media), or YES plates in which glucose was replaced with 3% glycerol or 3% glycerol + 0.1% glucose (gly + glu) or 2% galactose + 0.1% glucose (gal + glu). Plates photographed after 2-8 days at 32$^{o}$C. TTC: after 3 days' growth colonies on YES plate were overlaid with TTC-containing agarose and incubated for ~24 h to assess respiratory competence.

**B** DCHF-DA staining to assess levels of reactive oxygen species. Cells of the indicated strains were incubated in the ROS indicator DCFH-DA, which is converted to fluorescent DCF in the presence of ROS, and imaged under brightfield and 488 nm illumination. Scale bar, 10 $\mu$m.

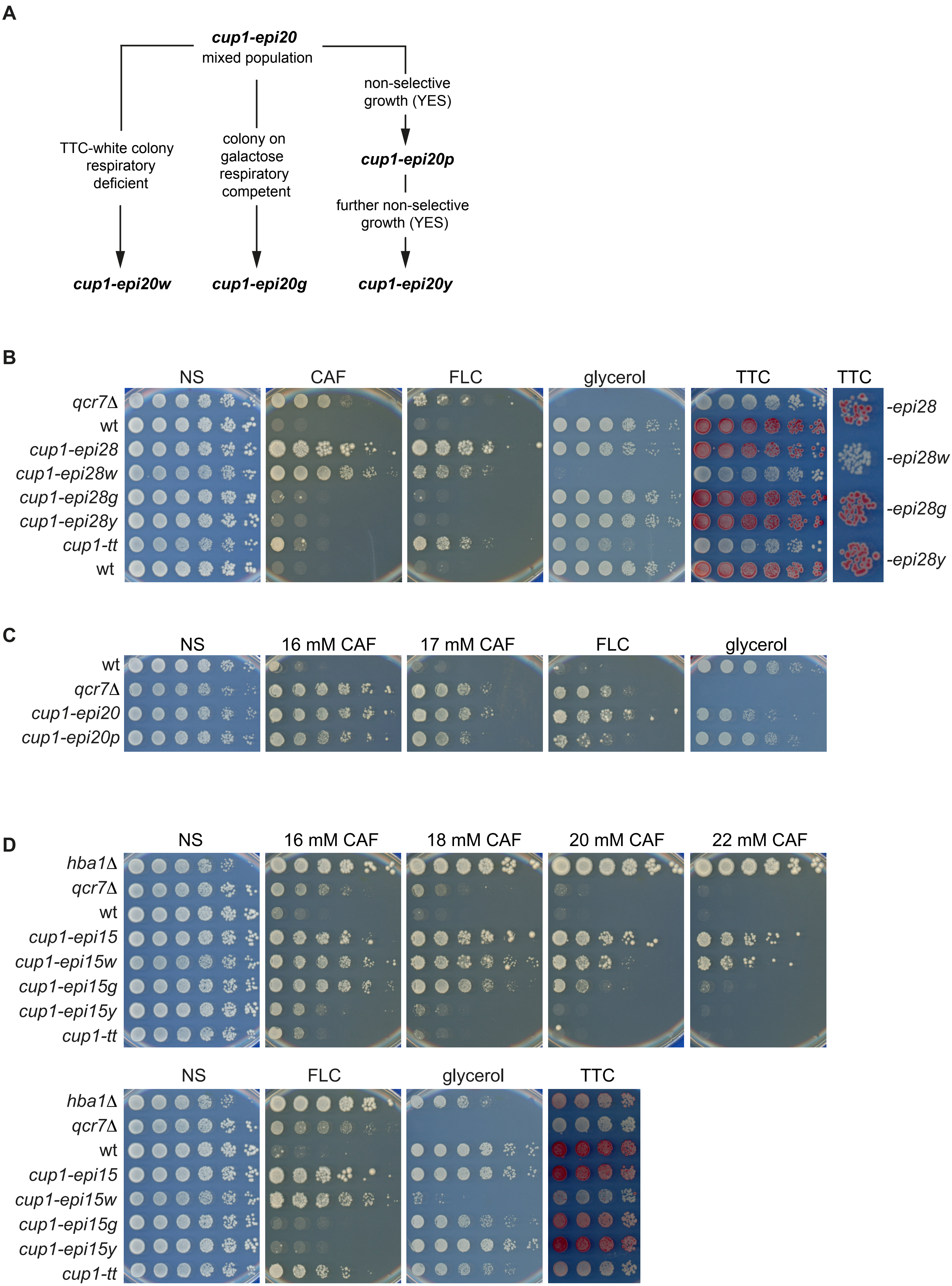


Fellas et al S7

**Figure S7: Phenotypes of *cup1* epimutant subpopulations**

**A** Isolation of *cup1-epi20* derivatives. *w* derivatives identified as TTC-white colonies; *g* derivatives selected for respiratory competence (growth on galactose + 0.1% glucose); *p* derivative isolated after extensive growth on non-selective YES media; *y* derivatives isolated after further growth on non-selective YES media; further details in Materials & Methods. **B-D** Growth assays to assess resistance to insults and respiratory competence of isolates derived from *cup1* epimutant mixed populations: Panels show five-fold serial dilutions of indicated isolates spotted onto non-selective plates (NS; YES media) or plates containing indicated concentrations of caffeine, 0.3 mM fluconazole (FLC), YES containing 3% glycerol rather than glucose, and TTC overlay to assay respiratory competence. **B**, *cup1-epi28* and derivatives; **C**, *cup1-epi20* and *cup1-epi20p* derivative; **D**, *cup1-epi15* and derivatives.

| strain number | relevant genotype | source |
|---|---|---|
| *972 h-* | *h-* | Lab stock |
| B4413 | *h- hba1-epi1, originally UR-1* | Torres-Garcia et al, 2020b |
| B4415 | *h- ppr4-epi3, originally UR-3* | Torres-Garcia et al, 2020b |
| B4416 | *h- UR-4* | Torres-Garcia et al, 2020b |
| B4417 | *h- UR-5* | Torres-Garcia et al, 2020b |
| B4418 | *h- UR-6* | Torres-Garcia et al, 2020b |
| B4427 | *h- cup1-epi 15, originally UR15* | Torres-Garcia et al, 2020b |
| B4432 | *h- cup1-epi 20, originally UR20* | Torres-Garcia et al, 2020b |
| B4440 | *h- cup1-epi 28, originally UR28* | Torres-Garcia et al, 2020b |
| B6766 (B5005) | *h- cup1-tt* | Torres-Garcia et al, 2020b |
| B5336 | *h- GFP-Pap1* | This study |
| B5422 | *h- fio1Δ* | This study |
| B5423 | *h- mbx2Δ* | This study |
| B6800 | *h- ppr4Δ* | This study |
| B5665 | *h+ GFP-pap1* | This study |
| B5671 | *h- hba1Δ::NAT GFP-pap1* | This study |
| B5671 | *h- GFP-Pap1 hba1Δ* | This study |
| B5676 | *h- GFP-Pap1 cup1-tt* | This study |
| B5676 | *h- GFP-Pap1 cup1-tt* | This study |
| B5680 | *h- GFP-Pap1 ppr4Δ* | This study |
| B5680 | *h- GFP-Pap1 ppr4Δ* | This study |
| B5684 | *h- pap1Δ* | This study |
| B5688 | *h- ppr4Δ pap1Δ* | This study |
| B5777 | *h- cup1-tt pap1Δ* | This study |
| B5969 | *h- fip1Δ* | This study |
| B6081 | *h- PX2:caf5pro-GFP (PX2 locus: chr2 between mrpl39 and str1)* | This study |
| B6081 | *h- PX2:caf5pro-GFP* | This study |
| B6088 | *h- obr1pro-GFP* | This study |
| B6126 | *h- fip1Δ fio1Δ* | This study |
| B6252 | *h- PX2:caf5pro-GFP hba1Δ* | This study |
| B6256 | *h- obr1pro-GFP hba1Δ* | This study |
| B6350 | *h- chr2 Pxlocus:caf5pro-GFP ppr4Δ* | This study |
| B6350 | *h- PX2:caf5pro-GFP ppr4Δ* | This study |
| B6354 | *h- obr1pro-GFP ppr4Δ* | This study |
| B6831 | *h- hba1Δ* | This study |
| B7055 | *h- SPBPC1348.10cΔ* | This study |
| B7058 | *h- eno102Δ* | This study |
| B7059 | *h- hsp3105Δ* | This study |
| B7061 | *h- rps1801Δ* | This study |
| B7062 | *h- region-UR6_1Δ* | This study |
| B7077 | *h- dal2Δ* | This study |

| | | |
|---|---|---|
| B7138 | *h- grt1Δ* | This study |
| B7187 | *h- mug129Δ* | This study |
| B7189 | *h- fyv7Δ* | This study |
| B7191 | *h- SPAC8C9.04Δ* | This study |
| B7192 | *h- dtd1Δ* | This study |
| B7193 | *h- cgs1Δ* | This study |
| B7194 | *h- rps5Δ* | This study |
| B7228 | *h- tad2 P292A* | This study |
| B7231 | *h- region-UR4_1Δ* | This study |
| B7235 | *h- region-UR4_3Δ* | This study |
| B7238 | *h- region-UR4_4Δ* | This study |
| B7275 | *h- region-UR4_2Δ* | This study |
| B7593 | *h+ pap1Δ::KAN prt1Δ::NAT bfr1Δ::HYG pmd1Δ::NAT caf5Δ::KAN mfs1Δ::NAT* | SAK72; Kawashima et al, 2012 via Ken Sawin lab |
| B7604 | *h- ndi1Δ::NAT* | This study |
| B7606 | *h- sdh7Δ::NAT* | This study |
| B7608 | *h- atp2Δ::NAT* | This study |
| B7614 | *h- cox4Δ::NAT* | This study |
| B7641 | *h- qcr7Δ::NAT* | This study |
| B7854 | *h- region-UR5-1Δ* | This study |
| B7904 | *h- region-UR5-2Δ* | This study |
| B8126 | *h- reb1Δ* | This study |
| B8128 | *h- rpm1Δ* | This study |
| B8246 | *h- PX2:caf5pro-GFP cup1-tt* | This study |
| B8250 | *h- obr1pro-GFP cup1-tt* | This study |
| B8288 | *h- GFP-Pap1 cox4Δ::NAT* | This study |
| B8291 | *h- GFP-Pap1 qcr7Δ::NAT* | This study |
| B8297 | *h- PX2:caf5pro-GFP qcr7Δ::NAT* | This study |
| B8312 | *h- obr1pro-GFP qcr7Δ::NAT* | This study |
| B8402 | *h- cup1-epi15w* | This study |
| B8404 | *h- cup1-epi20w* | This study |
| B8408 | *h- cup1-epi28w* | This study |
| B8509 | *h- cup1-epi15g* | This study |
| B8510 | *h- cup1-epi28g* | This study |
| B8513 | *h- cup1-epi20g* | This study |
| B8556 | *h- cup1-epi15y* | This study |
| B8558 | *h- cup1-epi20p* | This study |
| B8563 | *h- cup1-epi20y* | This study |
| B8569 | *h- cup1-epi28y* | This study |

**Supplementary Table 1: *S. pombe* strains used in this study**

| Name | Sequence | Purpose |
|---|---|---|
| HH1_1348.10_sgRNA_F | CTAGAGGTCTCGGACTGATCCTTATGGTTGG CCCCTGTTTCGAGACCCTTCC | Golden Gate cloning C1348.10cΔ-sgRNA-F |
| HH2_1348.10_sgRNA_R | GGAAGGGTCTCGAAACAGGGGCCAACCATA AGGATCAGTCCGAGACCTCTAG | Golden Gate cloning C1348.10cΔ-sgRNA-R |
| AF322_FsgRNA_cgs1Δ | CTAGAGGTCTCGGACTCGATATGGTTGCTG GCCCCGGTTTCGAGACCCTTCC | Golden Gate cloning cgs1Δ-sgRNA-F |
| AF323_RsgRNA_cgs1Δ | GGAAGGGTCTCGAAACCGGGGCCAGCAACC ATATCGAGTCCGAGACCTCTAG | Golden Gate cloning cgs1Δ-sgRNA-R |
| HH29_dal2_sgRNA_F | CTAGAGGTCTCGGACTTATTCGTTGAAACGG GTGCCGTTTCGAGACCCTTCC | Golden Gate cloning dal2Δ-sgRNA-F |
| HH30_dal2_sgRNA_R | GGAAGGGTCTCGAAACGGCACCCGTTTCAA CGAATAAGTCCGAGACCTCTAG | Golden Gate cloning dal2Δ-sgRNA-R |
| AF334_FsgRNA_dtd1Δ | CTAGAGGTCTCGGACTTCTTTATCTTGTCGC TTCCGGTTTCGAGACCCTTCC | Golden Gate cloning dtd1Δ-sgRNA-F |
| AF335_RsgRNA_dtd1Δ | GGAAGGGTCTCGAAACCGGAAGCGACAAGA TAAAGAAGTCCGAGACCTCTAG | Golden Gate cloning dtd1Δ-sgRNA-R |
| HH15_eno102_sgRNA_F | CTAGAGGTCTCGGACTCAATATCATTGGCCC TGCCTGTTTCGAGACCCTTCC | Golden Gate cloning eno102Δ-sgRNA-F |
| HH16_eno102_sgRNA_R | GGAAGGGTCTCGAAACAGGCAGGGCCAATG ATATTGAGTCCGAGACCTCTAG | Golden Gate cloning eno102Δ-sgRNA-R |
| AF114_FsgRNA_fioΔ | CTAGAGGTCTCGGACTGACATGTCCCAATAC CCCGAGTTTCGAGACCCTTCC | Golden Gate cloning fioΔ-sgRNA-F |
| AF115_RsgRNA_fioΔ | GGAAGGGTCTCGAAACTCGGGGTATTGGGA CATGTCAGTCCGAGACCTCTAG | Golden Gate cloning fioΔ-sgRNA-R |
| AF340_FsgRNA_fyv7Δ | CTAGAGGTCTCGGACTTAAAAAAGGGCTCA GGTGAAGTTTCGAGACCCTTCC | Golden Gate cloning fyv7Δ-sgRNA-F |
| AF341_RsgRNA_fyv7Δ | GGAAGGGTCTCGAAACTTCACCTGAGCCCT TTTTTAAGTCCGAGACCTCTAG | Golden Gate cloning fyv7Δ-sgRNA-R |
| HH48_V2grt1sgRNA_F | CTAGAGGTCTCGGACTTTCTATAATATCATCT GGGGGTTTCGAGACCCTTCC | Golden Gate cloning grt1Δ-sgRNA-F |
| HH49_V2grt1sgRNA_R | GGAAGGGTCTCGAAACCCCCAGATGATATTA TAGAAAGTCCGAGACCTCTAG | Golden Gate cloning grt1Δ-sgRNA-R |
| HH22_hsp3105_sgRNA_F | CTAGAGGTCTCGGACTTGTACGAACCGCCG TGTGACGTTTCGAGACCCTTCC | Golden Gate cloning hsp3105Δ-sgRNA-F |
| HH23_hsp3105_sgRNA_R | GGAAGGGTCTCGAAACGTCACACGGCGGTT CGTACAAGTCCGAGACCTCTAG | Golden Gate cloning hsp3105Δ-sgRNA-R |
| AF120_FsgRNA_mbx2Δ | CTAGAGGTCTCGGACTTGCGTTTGCCTACGA TGACCGTTTCGAGACCCTTCC | Golden Gate cloning mbx2Δ-sgRNA-F |
| AF121_RsgRNA_mbx2Δ | GGAAGGGTCTCGAAACGGTCATCGTAGGCA AACGCAAGTCCGAGACCTCTAG | Golden Gate cloning mbx2Δ-sgRNA-R |
| AF352_FsgRNA_mug129Δ | CTAGAGGTCTCGGACTAGATGATGGTACGT GTACGGGTTTCGAGACCCTTCC | Golden Gate cloning mug129Δ-sgRNA-F |
| AF353_RsgRNA_mug129Δ | GGAAGGGTCTCGAAACCCGTACACGTACCA TCATCTAGTCCGAGACCTCTAG | Golden Gate cloning mug129Δ-sgRNA-R |
| AF132_FsgRNA_pap1Δ | CTAGAGGTCTCGGACTAATGGTGAGGATGT GGCGGAGTTTCGAGACCCTTCC | Golden Gate cloning pap1Δ-sgRNA-F |
| AF133_RsgRNA_pap1Δ | GGAAGGGTCTCGAAACTCCGCCACATCCTC ACCATTAGTCCGAGACCTCTAG | Golden Gate cloning pap1Δ-sgRNA-R |
| AF126_FsgRNA_ppr4Δ | CTAGAGGTCTCGGACTTCGGACTTGAAGGG TCGGGTGTTTCGAGACCCTTCC | Golden Gate cloning ppr4Δ-sgRNA-F |
| AF127_RsgRNA_ppr4Δ | GGAAGGGTCTCGAAACACCCGACCCTTCAA GTCCGAAGTCCGAGACCTCTAG | Golden Gate cloning ppr4Δ-sgRNA-R |
| AF462_FsgRNA_reb1D | CTAGAGGTCTCGGACTGAACAATATTACGGC ACCTCGTTTCGAGACCCTTCC | Golden Gate cloning reb1Δ-sgRNA-F |
| AF463_RsgRNA_reb1D | GGAAGGGTCTCGAAACGAGGTGCCGTAATA TTGTTCAGTCCGAGACCTCTAG | Golden Gate cloning reb1Δ-sgRNA-R |
| AF468_FsgRNA_rpm1D | CTAGAGGTCTCGGACTGGGCTTCATGGCAG CCCCCGGTTTCGAGACCCTTCC | Golden Gate cloning rpm1Δ-sgRNA-F |
| AF469_RsgRNA_rpm1D | GGAAGGGTCTCGAAACCGGGGGCTGCCATG AAGCCCAGTCCGAGACCTCTAG | Golden Gate cloning rpm1Δ-sgRNA-R |
| HH36_rps1801_sgRNA_F | CTAGAGGTCTCGGACTGACGAAGACCACGG TGGGTTGTTTCGAGACCCTTCC | Golden Gate cloning rps1801Δ-sgRNA-F |
| HH37_rps1801_sgRNA_R | GGAAGGGTCTCGAAACAACCCACCGTGGTC TTCGTCAGTCCGAGACCTCTAG | Golden Gate cloning rps1801Δ-sgRNA-R |
| AF346_FsgRNA_rps5Δ | CTAGAGGTCTCGGACTCTCATCAAGAGAGA CGCCCGGTTTCGAGACCCTTCC | Golden Gate cloning rps5Δ-sgRNA-F |

| | | |
|---|---|---|
| AF347_RsgRNA_rps5Δ | GGAAGGGTCTCGAAACCGGGCGTCTCTCTT GATGAGAGTCCGAGACCTCTAG | Golden Gate cloning rps5Δ-sgRNA-R |
| WC407-sgi-pap1N-F | CTAGAGGTCTCGGACTTGACAACGTCTCAGT TTGTCGTTTCGAGACCCTTCC | Golden Gate cloning sgRNA pap1 N term |
| WC408-sgi-pap1N-R | GGAAGGGTCTCGAAACGACAAACTGAGACG TTGTCAAGTCCGAGACCTCTAG | Golden Gate cloning sgRNA pap1 N term |
| ST-1054-sgi-chrII | CTAGAGGTCTCGGACTAGGCCTTAATATTAA CCCCCGTTTCGAGACCCTTCC | Golden Gate cloning sgRNA PX2 locus |
| ST-1055-sgi-chrII | GGAAGGGTCTCGAAACGGGGGTTAATATTA AGGCCTAGTCCGAGACCTCTAG | Golden Gate cloning sgRNA PX2 locus |
| WC655-sg-F-PX-pro-ins | CTAGAGGTCTCGGACTAGAATATTCGCACTT AATAGGTTTCGAGACCCTTCC | Golden Gate cloning sgRNA PX2 locus, promoter insertion |
| WC656-sg-R-PX-pro-ins | GGAAGGGTCTCGAAACCTATTAAGTGCGAAT ATTCTAGTCCGAGACCTCTAG | Golden Gate cloning sgRNA PX2 locus, promoter insertion |
| AF328_FsgRNA_SPAC8C9.04Δ | CTAGAGGTCTCGGACTAGCCGACGTAGTAC CGGGACGTTTCGAGACCCTTCC | Golden Gate cloning SPAC8C9.04Δ-sgRNA-F |
| AF329_RsgRNA_SPAC8C9.04Δ | GGAAGGGTCTCGAAACGTCCCGGTACTACG TCGGCTAGTCCGAGACCTCTAG | Golden Gate cloning SPAC8C9.04Δ-sgRNA-R |
| HH50_tad2_sgRNA_F | CTAGAGGTCTCGGACTTTCAACAGTCACGTA TAAAGGTTTCGAGACCCTTCC | Golden Gate cloning tad2-sgRNA-F |
| HH51_tad2_sgRNA_R | GGAAGGGTCTCGAAACCTTTATACGTGACTG TTGAAAGTCCGAGACCTCTAG | Golden Gate cloning tad2-sgRNA-R |
| HH60_UR4_2_sgRNA_F | CTAGAGGTCTCGGACTTCATTACAGCACCCG CACGGGTTTCGAGACCCTTCC | Golden Gate cloning UR4_2-sgRNA-F |
| HH61_UR4_2_sgRNA_R | GGAAGGGTCTCGAAACCCGTGCGGGTGCTG TAATGAAGTCCGAGACCTCTAG | Golden Gate cloning UR4_2-sgRNA-R |
| WC657-PX-GFP-F | TAAGCTTGCTATGTTCTTTAGCTGCGTGTCT ATCATCTCTTCTTGTTTGGCTTGGCACGTGC CATTTTACCATATGAACAGTAAAGGAGAAGA ACTTTTC | insertion of GFP at PX2 locus |
| WC658-PX-GFP-adhT-R | TGCTTATAGGAGACAACGTGCTTAACTGAAA TGTTAACAGTTGCATACTTAGACTTATCTTGG ATAGTGAGTTTGGCCCGGTAGAGGTGTGGT CAATAAG | insertion of GFP at PX2 locus |
| HH3_1348.10_HR_F | GTACAATAAAGGTAAATCTATCTATTCTAAAT TTCTCACATGCCTGAGAATCCATTGAAACAA CATCAATGGAATCTTTTTGCAGAAAATAAAAA AAGTT | Making C1348.10cΔ-HR-template – F |
| HH4_1348.10_HR_R | AATATTCTTCCGTGTAGTACACTTTTCCATGC GAACTTCAATTTTTACAGCTTTTGTTTGAACT TTTTTTATTTTCTGCAAAAAGATTCCATTGAT GTTG | Making C1348.10cΔ-HR-template – R |
| AF324_HR_F_cgs1Δ | CCGCGAGATTGTGATTCACTGAGAGTAAGAA ACAAAAAACCGCATTTTTTGAAAGAAAGGCA CGTAAACTTTAGGAGACAAATTTCTATAATCT GTTTTC | Making cgs1Δ-HR-template – F |
| AF325_HR_R_cgs1Δ | AACAACAGGGAAATTCTTTAGCATATCATGA CAGGTCATTATTACAAAGATAGTTTCAAAGAA AACAGATTATAGAAATTTGTCTCCTAAAGTTT ACGTG | Making cgs1Δ-HR-template – R |
| HH31_dal2_HR_F | AAAATAATGTGCGAAACGAACCGGTAATCTT AGGTACATAAAATAAGACAGAAAAAAGAATA ATTAAAAAGTAAATCCCCTTTCGTAAAGAGAA AGTAAG | Making dal2Δ-HR-template – F |
| HH32_dal2_HR_R | GAGTCCCTTTGCTCTGGAACCTCTAATTTCC TTTTTCTTTCTGTTTATTAGCATTCCCATCTTA CTTTCTCTTTACGAAAGGGGATTTACTTTTTA ATTA | Making dal2Δ-HR-template – R |
| AF336_HR_F_dtd1Δ | ATATTATTGTTGTTATTATTAATAGTCTTCGG TTCTGTCCATCCTTCGTTACCTTTTTTTTTTT GAACAGTAGAAAACATTAGATTTATTAAATGA TAT | Making dtd1Δ-HR-template – F |
| AF337_HR_R_dtd1Δ | ATAAAGATTACTGAATTTTAATTTCCAAAATC GTCAACTAGTTCTAATGGTTGAGTACGGATA TCATTTAATAAATCTAATGTTTTCTACTGTTCA AAAA | Making dtd1Δ-HR-template – R |
| HH17_eno102_HR_F | CAAAAGATAATTCCATATATAACCGTAAAACG GTAAATACAGCATCAATGAAAATATAATAGCA | Making eno102Δ-HR-template – F |

| | | |
|---|---|---|
| | AAAAATTATCGCAACTTTTTAATATAAAAAG TTGT | |
| HH18_eno102_HR_R | GTATATATATATATATTTATACCTACTATCCAA ATCACAGACTTCAGAAGAATTTTGAAAACAA CTTTTTTATATTAAAAAGTTGCGATAATTTTTT GCT | Making eno102Δ-HR-template – R |
| AF116_HR_F_fioΔ | TGTTTTCATTTCTTTTTTTTCTGTTCTTTTTTTT TTTTAATATCTTTCCTTATTCTTCTTTTCTCTC TTGCGCGCTATAAAAATAAGAGATGAAAGAA AA | Making fioΔ-HR-template – F |
| AF117_HR_R_fioΔ | TAAAACGGAGTAAAAAAAAAAGACACACCAA CTGTAATTTTGGGGGTAGGAAACACATATTT TTCTTTCATCTCTTATTTTTATAGCGCGCAAG AGAGAA | Making fioΔ-HR-template – R |
| AF342_HR_F_fyv7Δ | TCAGGAAATACATTGGATTTTGGTTATTTCTT TATTAAGGTATTTGATTTGAGATCGAAGAATT CTGGGATCAATTAGAAGCATGTATGTTTTAG TGAAG | Making fyv7Δ-HR-template – F |
| AF343_HR_R_fyv7Δ | CCCAATTAATAAAGATTTCAATTTTTGTGTGC GTGTTGTATGGCAGAACATCAGAGATTGCTT CACTAAAACATACATGCTTCTAATTGATCCCA GAATT | Making fyv7Δ-HR-template – R |
| HH10_grt1_HR_F | CCAACTAAGTGGGTGGATTATTTGATCAAAG CCAAATGGATTTAATCTAAACCTTTTGTGAAT CTTATATAGCTTTCGTTTGTGACACGTTGGG AATAAA | Making grt1Δ-HR-template – F |
| HH11_grt1_HR_R | AATTTTTGTGAGTTTATTCTTATATTTTCAAAA AAAGCTATTAAAAACGGCCCTCTGCATTTTAT TCCCAACGTGTCACAAACGAAAGCTATATAA GATT | Making grt1Δ-HR-template – R |
| HH24_hsp3105_HR_F | TTTTTAAAATATAACGAGGATATTTTAACTGA TAACCCAAATTTGCGACATTGAATTGTATTGA AAATTGAATTAAAAAATCAAAATTTCTTTTTTG GAA | Making hsp3105Δ-HR-template – F |
| HH25_hsp3105_HR_R | TGCCATTTCAAACATTAGAAAAAATAAAAAAA ATGTGAGAATCCCAAAACACGATCGGTTTTC CAAAAAAGAAATTTTGATTTTTTAATTCAATTT TCAA | Making hsp3105Δ-HR-template – R |
| AF122_HR_F_mbx2Δ | ATACCTGAATTTTGGTTAACCTCGTCGACTC TTTTTCTCTTTTCTCGTCAAACCTTACTAAAT ATATACAACGTCTAATAAGAATATCATAGAAG TCAAT | Making mbx2Δ-HR-template – F |
| AF123_HR_R_mbx2Δ | TACTTGTATACGATTCATTAGATCCAGGGAG GACACATGATGAAGACATTTGAAAGGGAGAT TGACTTCTATGATATTCTTATTAGACGTTGTA TATATT | Making mbx2Δ-HR-template – R |
| AF354_HR_F_mug129Δ | AGTTTGTAGACTCGAAAAAAAAACCACATTG TTTTTTCAAATAGTATTGTCAAAAGATACTTC GTGAAGGAGAAGTACATCACAAGTTTGATTA TTTGGA | Making mug129Δ-HR-template – F |
| AF355_HR_R_mug129Δ | TTATTGTTGATTTTTTATATCTCGCGCGGTCG TTTTGTTTTCCAATTTTAAACTTATCTGTCCAA ATAATCAAACTTGTGATGTACTTCTCCTTCAC GAA | Making mug129Δ-HR-template – R |
| AF134_HR_F_pap1Δ | TATTAATTAAAACAATGGAAATACATATAAAA ATAAGCAAAAAGTCGTTATAACATCGATATA GATCAAACCAAGCAGGTAATTATCAATTAATT AACAC | Making pap1Δ-HR-template – F |
| AF135_HR_R_pap1Δ | CTCTTATTTTATAAATTTTTTATCCTATCGATA CTACATTTTTCAAAGGAGCATACATTCGTGTT AATTAATTGATAATTACCTGCTTGGTTTGATC TAT | Making pap1Δ-HR-template – R |
| AF128_HR_F_ppr4Δ | CACCCAGAGCTAAGTTTACAAGTCCAATTTT TTGACAAAGGCAGCATACGGCGAGACGGCT GAGTTCACAATTGAAATATGTAATAACCTGAA ATTAGAC | Making ppr4Δ-HR-template – F |

| | | |
|---|---|---|
| AF129_HR_R_ppr4Δ | ATTTATTTTTCTCTTTTCCTCATAATCACTTTATCTTTGTAATTTTTTGACTTTCACAGAGTCTAATTTCAGGTTATTACATATTTCAATTGTGAACTCA | Making ppr4Δ-HR-template – R |
| AF464_HR_F_reb1D | TAATAAGGACTTGTAAAAAGATACAAAGTTACTTAAGTAAGTTAAAACAAATTTTTTCGTTCAAAAAATTGTAAGGACGTAAAGATTATATTTTTGAAGT | Making reb1Δ-HR-template – F |
| AF465_HR_R_reb1D | CACGAAGCGTTTTGTACGATATTAGCGATTGATAAGTTGAAGTGATTACTCAATTATAGTACTTCAAAAATATAATCTTTACGTCCTTACAATTTTTTGA | Making reb1Δ-HR-template – R |
| AF470_HR_F_rpm1D | AAAGTGCTCTGGTTCAGCTCTCTAGTCGAATTTACCATCAACACTATTTTTGGTGTTACTAATTATAGGGTTTTTGCCCACAAACTTATTAAAACTATTT | Making rpm1Δ-HR-template – F |
| AF471_HR_R_rpm1D | TCTCGGTAAATTGATAAAATGCAGGATTGAGTATTTAATTAAGTAAAATTATATCCATAAAAATAGTTTTAATAAGTTTGTGGGCAAAAACCCTATAATT | Making rpm1Δ-HR-template – R |
| HH38_rps1801_HR_F | AAGAATCAGAAAACTATAATTTGCAAGAATAATGGAAATTTATTATCCAAGTAATATTAGGCTAATCATTTTACTTATTTTTTCCGGAAGCTGCTGGTAA | Making rps1801Δ-HR-template – F |
| HH39_rps1801_HR_R | TGACTGCTTTTTGAGAGTGCTTCAGTGTTTGGGCTTGTTCTAGTTGATATACTTTTGTACTTACCAGCAGCTTCCGGAAAAAATAAGTAAAATGATTAGC | Making rps1801Δ-HR-template – R |
| AF348_HR_F_rps5Δ | GGTAGGGTTTACGAACAGTCACAGTTTGTTTCCATAAGCTAAAGAGCTTGTATGAACTGAGAAAAAACCACAACGCAAGAAGACTTATTTGTATTAATGG | Making rps5Δ-HR-template – F |
| AF349_HR_R_rps5Δ | CATTGCCTTTTACTACGAGAAGTTAAAGCACAAAGATTAAATGCAAGGGAGTAACAAAATCCATTAATACAAATAAGTCTTCTTGCGTTGTGGTTTTTTC | Making rps5Δ-HR-template – R |
| AF330_HR_F_SPAC8C9.04Δ | ACGCCAAGATTTACAACTTCTACCAACCACCAAACAAGAACGCAGAGCGTGTAAAAGTAGAACAAAAATTGAGTTGAATAGGTTTACCTGGAAGCATTGG | Making SPAC8C9.04Δ-HR-template – F |
| AF331_HR_R_SPAC8C9.04Δ | TGTCGCTAAAAGGTGAATTTTTCAACATACAAAAATAAAAGACGGAAATCAAACATATGCCCAATGCTTCCAGGTAAACCTATTCAACTCAATTTTTGTT | Making SPAC8C9.04Δ-HR-template – R |
| HH52_tad2P292A_HR_Flong | AGGGGATTCGACATGCTGAACTTATCGCTATAGAAAAAATTTTAGAACACTACCCAGCATCTGTTTTCAAAGAAACAACTTTATACGTGACTGTTGAAGCGTGTCTAATGTGTGCTGCTG | Making tad22P292A-HR-template – F |
| HH53_tad2P292A_HR_R | TCTCGACCTTCGTGTTCGTG | Making tad22P292A-HR-template – R |
| HH56_UR4_1_HR_F | GAACCGTATATTAATGCAAACTAAACCAACTTCGTTATATAGGAATTGCGATAAAAAAAGTTTGTAAGTATTACAAGATATTCACATCATAATGTTTTTCCCAAATAAATAACAATGCTT | Making UR4_1-HR-template – F |
| HH57_UR4_1_HR_R | TAACATTTGTTACTCAATAGAAATTTAAGGTAACATTGATCATAAAGTAAAAGCTTGCCGTACTAAATTGTATCCGCCTAACGTTTGCGAATTTTGCAGCAAGCATTGTTATTTATTTGG | Making UR4_1-HR-template – R |
| HH62_UR4_2_HR_F | TTATTTCTTTCATATATATAGATATAATCTGCCATTTTTAGATTCATCAAGAACACAATAATCTTCAAGTATTTAGATTTTTGCATTCGCTGGATAAACAAATAGAATTCGTCAATCGAC | Making UR4_2-HR-template – F |
| HH63_UR4_2_HR_R | TTGGTCACACTGTTTTGGACTATCGATACAAAGTGCGATGTTCGCTATTCTTAAAAAACCTTATAAAGACAGTAAAGTAGGTACCAATCTTTAATATTTAGTCGATTGACGAATTCTATT | Making UR4_2-HR-template – R |
| HH66_UR4_3_HR_F | ATAGAATTCGTCAATCGACTAAATATTAAAGATTGGTACCTACTTTACTGTCTTTATAAGGTTT | Making UR4_3-HR-template – F |

| | | |
|---|---|---|
| | TTTAAGAATAGCGAACATCGCACTTTGTATC GATAGATCCTTTATGTAACTTTATC | |
| HH67_UR4_3_HR_R | ATATTAGTAGTGTGCAGATCGCAAGAAGAAG TTACCGTAGTTTACCCAATCATTATAGCACAT AGCAAACTGCGTAGCAAACATATAAAGTCAC GTAGCTGATAAAGTTACATAAAGGAT | Making UR4_3-HR-template – R |
| HH70_UR4_4_HR_F | AGTTGCTGATAAATCGTTCACAGAAAATAGA TTGCTAAGCACTGAATAGCAGCATTCCCTTA CGAAATCAATCAAGATTAAACTTTAACTTTCT AATACCCAGAAAATGTAGAGAATAAG | Making UR4_4-HR-template – F |
| HH71_UR4_4_HR_R | AAATTACATTTCGCAGGAAGTTTAGATAATTG TAAATGTGTTAAGGAGTATATAGTAAGAATG CAAGAAATAAATCAAGGACGTAAAATACTGC CTTCCTCTTATTCTCTACATTTTCTG | Making UR4_4-HR-template – R |
| HH80_UR5_1_HR_F | ATTTTCAATAACTGAAGTACGATGTTTATAGC AATATCATTATTAGGAGAAAACTTGATATGGA AATACTTTATGAAATAGTTGCTTCAATATTAC ATTAGGAAATGAAATGGATGCGCA | Making UR5_1-HR-template – F |
| HH81_UR5_1_HR_R | GCACACAGAGCATTTCCGAGGTCAACCTATT TAGTTTGCCTTGGTAATCGCCAATTGTATCA AGTTTCCAACTCCTTGCAGACGGATGCAAAT TGGACGGTGCGCATCCATTTCATTTCC | Making UR5_1-HR-template – R |
| HH84_UR5_2_HR_F | TGTTCAAATCCTGCAATACCAATCTCTTCCAT AGAAGGAAGACAATGGGCGAATTACCACAC AATCTTCTAATGTATTATTTCTAGCAAAAAAA GCAAGAGTAGCAGAATGCCGAAATTA | Making UR5_2-HR-template – F |
| HH85_UR5_2_HR_R | TAATTTCATGGCATTAGTACCGTGGGCTAGT TTCGTAATGCATTTAACTCATGACTTTTAAGT TTATAAACTATCAGTTGTTACGGATACTTTAG TATGTTAATTTCGGCATTCTGCTAC | Making UR5_2-HR-template – R |
| WC443-GFP-pap1-F | CTTATTTTATAAATTTTTTATCCTATCGATACT ACATTTTTCAAAGGAGCATACATTCGTGTTAA TTAATTGATAATTATGAACAGTAAAGGAGAA GAAC | N-terminal tagging of pap1, SpEDIT |
| WC444-GFP-pap1-R | AATCGGCGGATTGCTCTGGTTCCGCCTTTGC AATTGGAATGTTTGAAGTAGATGACAACGTC TCAGTTTGTCCTGACATTTTGTATAGTTCATC CATGCC | N-terminal tagging of pap1, SpEDIT |
| WD77-atp2-Bah-KO-F | ATGTATACTCAAAAAAGTGGGAATCATTCCA TAGTCCTTTTCTACGTAGCCAACCTAACCAA CCCTTTGCATACGATCGGATATACCAACCGC AATCAAACGGATCCCCGGGTTAATTAA | gene deletion, Bahler method |
| WD78-atp2-Bah-KO-R | CCAAGCGGTAAGCAAATAACCGCTGAAGAG AATTTCTGGCTTGAACAAGATTGATGACAAG TTTTGTGCAAAAAACATCAAAAAGTCTTAGCA AATTGATGAATTCGAGCTCGTTTAAAC | gene deletion, Bahler method |
| WD79-cox4-Bah-KO-F | ACCAAATTTCGTTACACAATACCGGACCAAC GTTGGAAACTACACAACACCCCATCTCACCC ATTATCAGGTTATAAATCATCAATAATTGTCT TTTAAGCGGATCCCCGGGTTAATTAA | gene deletion, Bahler method |
| WD80-cox4-Bah-KO-R | CATACAGTTTTCATTGATTTTTCAAAGTTCAA ATAAAAGAGCTAAATTGTTATAAAAGAGTATG AAAATAGCAAGGGGATATGGAAGATTGGTAA GCATAGAATTCGAGCTCGTTTAAAC | gene deletion, Bahler method |
| WD83-ndi1-Bah-KO-F | ATTTGTCATTATCTCATACATTTTTTTACTCAC GCCTCCCTACCACTTATCACTTAGTCCTATTT AAGGGGAAATTTGTTTACTTCTTTTTTCTTAA AAACGGATCCCCGGGTTAATTAA | gene deletion, Bahler method |
| WD84-ndi1-Bah-KO-R | AAATACGTTCCTTTTTCAAGTATAAAATAATC ATCTTATGATTAACGACAAAGCACAAGGAAA CAATTGCTTGTAAACCAAAAGTATTTCATTCC GTTCAGAATTCGAGCTCGTTTAAAC | gene deletion, Bahler method |
| WD85-qcr7-Bah-KO-F | GATCTTACCTTCTTCCAATATTCAAAACTAAG AGTTTTATTGATTTCAGTGCAAATTTTGTAGT GACGAAATTTTTTTTTGCTGAAGTGATTGTTT AAGTCGGATCCCCGGGTTAATTAA | gene deletion, Bahler method |

| | | |
|---|---|---|
| WD86-qcr7-Bah-KO-R | ACTTAGAAAATAAAATTCTCACATCCGTTTACTACTCTTTTAATTTTCTTGTCGGTTTTCATGAGAGGAATAAGTGGATTAAAGGAGGACATGCAAACTTGAATTCGAGCTCGTTTAAAC | gene deletion, Bahler method |
| WD89-sdh7-Bah-KO-F | GCGATGAATAGCAAACTGCCTTCCACATCACAACCACAACAAGAAACTTAAAAGGAATTCAATTTTTGAAAAGACGGTATCTAAAGAAAATCATATACCACGGATCCCCGGGTTAATTAA | gene deletion, Bahler method |
| WD90-sdh7-Bah-KO-R | GATGTCGTCTCAAAGTGCCAATAAAAATGACGACAACTCAAACAATCAAAATCCTTCTAATAACAAAGAGATGTCTGATAAAGAATCAGATTCCGCGACCGAATTCGAGCTCGTTTAAAC | gene deletion, Bahler method |
| WC665-PX-GFP-caf5-pro-F | TAACTATCTTTTAAAGAGGATTTGACATTGTACTAAGGGCTGAGACAACTCGCATTAAATAAGCAAAGGCTGAATTAAGTTAAGTAACTGTTGTATTCGC | creation of caf5 promoter GFP reporter |
| WC666-PX-GFP-caf5-pro-R | TGCCCATTAACATCACCATCTAATTCAACAAGAATTGGGACAACTCCAGTGAAAAGTTCTTCTCCTTTACTGTTCATTTAAATGACCGTATGTAAAAACC | creation of caf5 promoter GFP reporter |
| WC663-PX-GFP-obr1-pro-F | TAACTATCTTTTAAAGAGGATTTGACATTGTACTAAGGGCTGAGACAACTCGCATTAAATAAGCAAAGGCTGAATTAAGAACAGTGGCGCAGCTACATCG | creation of obr1 promoter GFP reporter |
| WC664-PX-GFP-obr1-pro-R | TGTGCCCATTAACATCACCATCTAATTCAACAAGAATTGGGACAACTCCAGTGAAAAGTTCTTCTCCTTTACTGTTCATACTGGGTTTAATAAAACGCTG | creation of obr1 promoter GFP reporter |
| WB9_act1_F | CCCAAATCCAACCGTGAGAAGATG | qPCR act1 negative control locus |
| WB10_act1_R | CCAGAGTCCAAGACGATACCAGTG | qPCR act1 negative control locus |
| WC613_caf5_F | TGAGGTTTTTGGCCGGTTTC | qPCR caf5 ORF |
| WC614_caf5_R | AAGCAACGGTCCCAAAACTG | qPCR caf5 ORF |
| WD629-obr1-up-qF | ACTCATCAAGCGAGCCAAAG | qPCR obr1 promoter region |
| WD630-obr1-up-qR | TCCACAGGTGTTGGCATTTG | qPCR obr1 promoter region |
| WD637-srx1-up-qF | AAGAACACAAACCCAGACCTCG | qPCR srx1 promoter region |
| WD638-srx1-up-qR | GTCAGCATGTTAGCGAAACAC | qPCR srx1 promoter region |
| WD647-bfr1-up-qF | TTGCTCCTCGGTGTTTAGTCG | qPCR bfr1 promoter region |
| WD648-bfr1-up-qR | GCCAAACGCTAGGAAGATGATG | qPCR bfr1 promoter region |
| WD203-caf5-up-F | AGCAACAAGATGATTTTGCAC | qPCR caf5 promoter region |
| WD204-caf5-up-R | TCCTAATATACCCTTCGTGCC | qPCR caf5 promoter region |

**Supplementary Table 2: Oligonucleotides used in this study**
